# Reproductive innovation enabled radiation in the deep sea during an ecological crisis

**DOI:** 10.1101/2024.01.12.575380

**Authors:** Chase D. Brownstein, Katerina L. Zapfe, Spencer Lott, Richard Harrington, Ava Ghezelayagh, Alex Dornburg, Thomas J. Near

## Abstract

Major ecological transitions are thought to fuel evolutionary radiations, but whether they are contingent on the evolution of certain traits is unclear. We show that the rapid ecological transition of anglerfishes into pelagic habitats during a period of major global warming coincided with the origins of sexual parasitism, in which male anglerfishes temporarily attach or permanently fuse to females to mate. A phylogenomic reconstruction of the evolutionary history of anglerfishes provides a strong inference for the convergent evolution of permanently-fusing deep-sea anglerfishes and their degenerate immune genes. Our results support that sexual parasitism was enabled by the degeneration of adaptive immunity and ancestral sexual size dimorphism. The combination of these traits facilitated the transition of pelagic anglerfishes into novel ecologies available in the deep open oceans after evolving from benthic ancestors. These results show how seemingly unrelated physiological and reproductive traits interact synergistically to drive evolutionary radiation in novel environments.

## Introduction

The origin of biodiversity can lie in the interplay of selective pressures with constraints imposed on form and function ^1–6^. A classic example of this phenomenon is the evolution of traits called key innovations that allow a lineage to utilize new habitats or resources ^7^. Key innovations are routinely invoked to explain how lineages are able to rapidly exploit ecological opportunities ^7–9^. Despite their potential importance to understanding the generation of major portions of biodiversity and the origin of novel phenotypes, investigations of key innovation almost exclusively focus on single traits rather than considering combinations of multiple features that collectively produce effects of interest ^10^. A focus on synergistic trait combinations, or synnovations, could prove useful for understanding how modifications in biological systems essential for survival, which are often characterized by coincident interactions among several traits, have influenced evolutionary history of hyperdiverse lineages ^10^. Understanding how synergistic traits arise simultaneously or predate modifications to these key biological systems would provide the historical context to develop general rules about the dynamics of biological diversification ^9,11,12^.

Immunity, which allows the body to defend itself from foreign biological pathogens, is a classic example of a biological system whose evolution is theoretically constrained by its importance ^13–17^. Vertebrates are unique in encoding numerous families of receptors that differentiate normal cells (self) from pathogens and infected cells (non-self). When an endogenous (“self”) antigen is presented to an immune cell, the cell is not engaged and there is no immune response. In contrast, when a pathogen-derived peptide (“non-self”) is presented, the immune cell can be engaged, resulting in destruction of the infected cell. This presentation of self and (if a cell is infected) non-self antigens is orchestrated by major histocompatibility complex (MHC) molecules in concert with a suite of associated genes and has become a pivotal aspect of vertebrate pathogen defense and immune memory. However, several species of ray-finned fishes, including seahorses that possess male pregnancy, have lost or degenerated the genetic basis of this physiological system ^18–21^.

Anglerfishes (Lophioidei) are a clade of ray-finned fishes that occupy a variety of benthic and open water marine habitats. The anglerfish clade Ceratioidea are the most species-rich lineage of vertebrates found in the bathypelagic, or ‘midnight,’ zone of the deep sea ^22^. Ceratioid anglerfishes have become well known for their reproductive habits, in which males act as sexual parasites and temporarily attach, permanently fuse to females, or facultatively switch from both of these modes to mate ^23,24^. This unusual reproductive mode is thought to increase the probability of successful reproduction once a mate has been found in the midnight zone, Earth’s largest and most homogenous habitat ^22–24^. In addition to compromising the MHC, sexual parasitism in several lineages of anglerfishes appears linked to a degenerated suite of molecular receptors associated with losses to critical aspects of the adaptive immune response such as V(D)J recombination, the ability to generate high affinity antibodies, T and B cell development, and antigen display. Collectively, this degeneration of the adaptive immune response is hypothesized to have facilitated the evolution of the reproductive modes of ceratioid anglerfishes ^13^, but the evolution of these modes of sexual parasitism and the corresponding genomic changes remains unclear ^25^. Whether synergetic interactions have occurred between these molecular changes and other key aspects of the ceratoid anglerfish radiation, including extreme sexual dimorphism in body size and a shift in locomotor mode from benthic walking to open-water swimming, remains unclear, as does the confluence of these factors with major changes to oceanic conditions that occurred throughout the Cenozoic.

Here, we reconstruct the evolutionary history of anglerfishes using a phylogeny inferred from 975 ultraconserved elements and calibrated using the fossil record of anglerfishes and the closely related pufferfishes (Tetraodontoidei) ^26^. We show that the invasion of the midnight zone by anglerfishes from a benthic ancestor occurred rapidly during the Paleocene-Eocene Thermal Maximum, a period of major climate change. Our analyses demonstrate that key aspects of the peculiar reproductive mode of ceratioids, including degeneration of the genomic basis of the adaptive immune response and extreme sexual dimorphism, were present among anglerfishes prior to their evolutionary transition to the bathypelagic zone. We find no pattern in the reconstructed evolutionary history indicating that sexual parasitism directly spurred ceratioid diversification. Instead, sexual parasitism appears to have indirectly facilitated the radiation of anglerfishes by aiding the transition to a new habitat in which numerous ecological opportunities were available. Our results show how combinations of traits from disparate biological systems can spur adaptive radiation by allowing a clade to access new habitats and ecologies.

## Results

### Genomic discordance stifles resolution of anglerfish evolutionary history

Phylogenetic relationships among the major lineages of anglerfishes have evaded resolution even in the era of widespread genome sequencing ^13,27–31^. We inferred a time-calibrated phylogenomic tree of anglerfishes using a dataset of 975 ultraconserved element (UCE) loci sequenced for 244 individuals, including 222 specimens of anglerfishes (Lophioidei) and 20 species of their sister lineage, the pufferfishes, ocean sunfishes, and triggerfishes (Tetraodontoidei). We employed a multistep bioinformatic pipeline ^32^ (see Methods) to thoroughly scrutinize the sequence data. Using several tools (phyluce ^32^, CIAlign ^33^, IQ-TREE^34,35^, TOAST ^36^), we aligned and preprocessed UCE sequences, removed chimeric UCE sequence data, flagged sequences that violated common molecular evolutionary model assumptions, and removed aberrant gene trees generated from an initial run of UCE alignments in the likelihood-based phylogenetic software IQTREE 2. We created species trees based on the multispecies coalescent as implemented in ASTRAL-III ^37^ and used IQ-TREE 2 ^34^ to infer maximum likelihood phylogenies from partitioned and concatenated versions of our UCE dataset. Our phylogenomic methods aimed to minimize the effects of hidden paralogy, alignment quality, and model violations that are known to contribute to systematic error in phylogenetic inferences ^38^.

These UCE-inferred phylogenies resolve the deepest divergences in anglerfishes with strong support: Lophiidae (goosefishes) is the sister lineage of all other anglerfishes, batfishes (Ogcocephalidae) and the frogfishes and handfishes (Antennariidae) comprise a clade of benthic species with modified fins for walking along the seafloor ^39^, and the seafloor-walking sea toads (Chaunacidae) are the sister lineage of the deep-sea anglerfishes (Ceratioidea) (Figure 1; Figure S1-S8). Our UCE phylogeny is comparable to the results of recent phylogenomic studies of anglerfishes ^27,28^ but differs considerably from efforts to resolve their relationships using smaller nuclear and mitochondrial datasets ^29,40,41^ or morphological characters ^42^. Our strong inference of ceratioid monophyly (Figure 1) suggests that this lineage transitioned to open water habitats from an ancestor that used its pelvic fins to walk along the seafloor ^39^ in a manner reminiscent of major transitions at the origin of groups like whales and marine reptiles ^43–45^.

**Figure 1.**
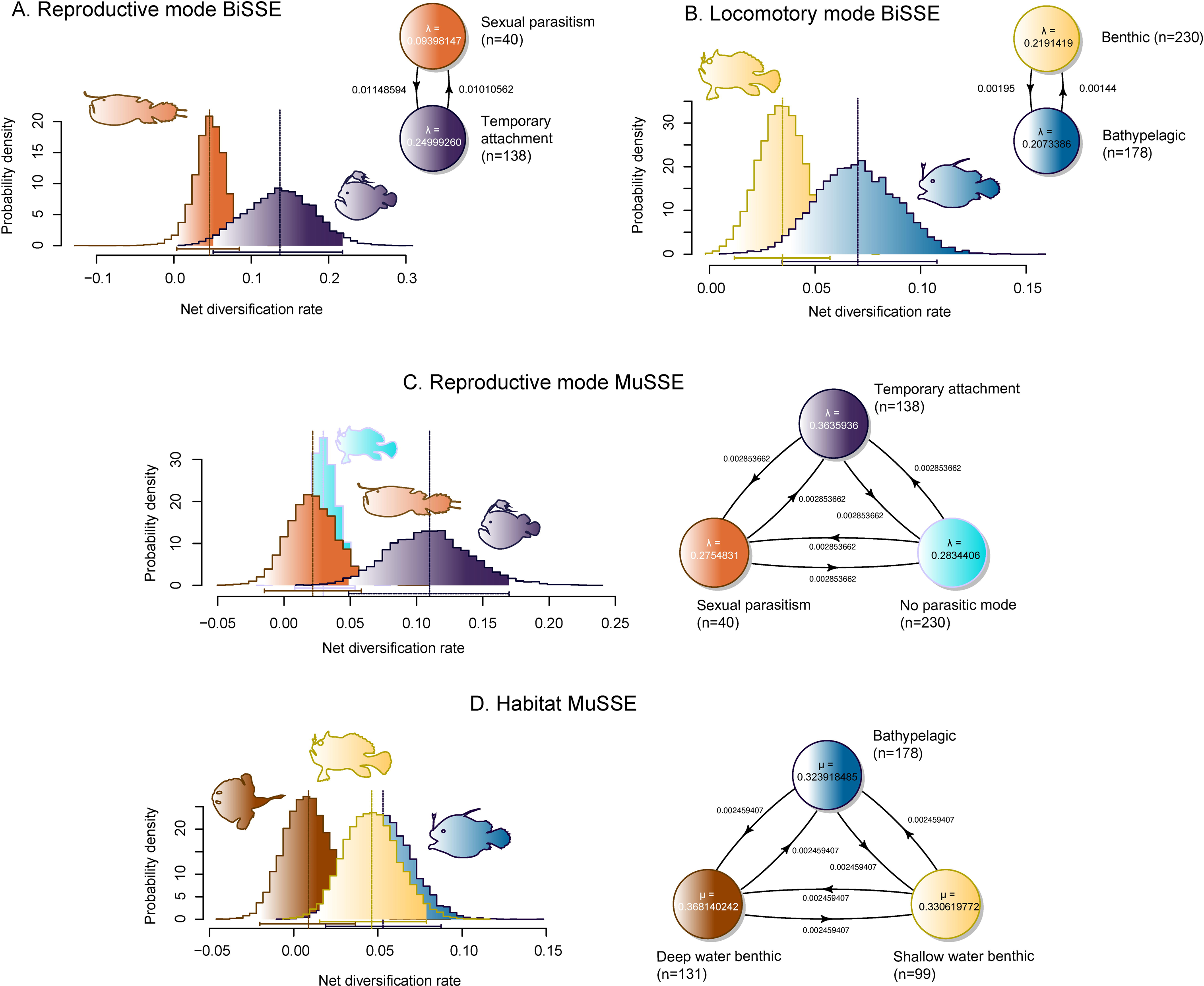
Anglerfishes invaded and radiated in the deep sea during a global warming event. Figure shows the Bayesian node-dated phylogeny produced from analysis of the 15 largest UCE partitions and 11 fossil calibrations in BEAST 2.6.6. Posterior tree sets generated from partitions that converged when analyzed were pooled and used to calibrate the target partitioned topology from IQTREE. Nodes are placed at median estimated ages, and gray bars at nodes indicate 95% highest posterior density intervals. Clear bars indicate bootstrap support values of less than 100 in the maximum likelihood tree resulting from analysis of the partitioned UCE dataset in IQTREE. Deep sea temperature curve is from Meckler et al. (2021). Fish illustrations by Julie Johnson.

Although we resolve of the relationships of major lineages of anglerfishes, we find that a high degree of uncertainty underlies the relationships of major lineages of deep-sea ceratioids: the obligate sexual parasitic Linophrynidae, *Neoceratias*, Ceratiidae, and *Centrophryne*, and the temporary attachers and facultative parasites in Gigantactinidae, *Thaumatichthys*, *Himantolophus*, Oneirodidae, Diceratiidae and *Melanocetus* (Figure 1; Figure S5; Figure S9). The high magnitude of phylogenetic discordance among individual loci indicates the backbone of the ceratioid phylogeny is in the so-called anomaly zone (Figure 1, Figure S9, Figure S10) ^38,46,47^, where variables such as large effective population size and rapid lineage divergence preclude adequate sorting of ancestral genetic variation and results in extensive incongruence among gene trees and the species tree. These results suggest a rapid pace of lineage divergence underlying early ceratioid evolution. While there is uncertainty among the major lineages of ceratioids, there is strong phylogenetic signal in the resolution of their monophyly and placement in the phylogeny of anglerfishes (Figure 1; Figure S3, Figure S9, Figure S10).

### Anglerfishes radiated during a period of extreme oceanic warming

We time-calibrated the anglerfish phylogenomic tree using a node-dating Bayesian method ^48^ with a partitioned UCE sequence dataset and a set of 13 fossil calibrations (Figure 1). The ages estimated for the major anglerfish lineages, including an earliest Paleogene age for crown Ceratioidea ^28^, are younger than previous estimates based on smaller phylogenetic datasets^29^. The median age of the most recent common ancestor (MRCA) of living anglerfishes is 87.7 Ma (95% highest posterior density: 71.7, 106.0 Ma). All major living lineages of anglerfishes originated in the Cenozoic (Figure 1). The posterior median age of the MRCA of Lophiidae is 55.6 Ma (95% HPD: 46.5, 80.8 Ma), the Antennariidae is 60.3 Ma (95% HPD: 52.1, 69.6 Ma), and the Ogcocephalidae is 30.2 Ma (95% HPD: 19.3, 44.1 Ma). Sea toads (Chaunacidae) represent a recent (Figure S11) diversification with an MRCA age of 6.0 Ma (95% HPD: 2.8, 10.3 Ma). The common ancestry of ceratioid anglerfishes lies in the Paleocene-Eocene, 51.8 Ma (95% HPD: 37.7, 67.4 Ma), in the relaxed clock analysis. Nearly all ceratioid family-level clades diverged 50 to 30 million years ago (Figure 1) during a period of high global temperatures called the Paleocene-Eocene Thermal Maximum (PETM)^49–53^, which induced extinction throughout the world’s oceans ^49,54^. The origination of ceratioid clades (Figure 1) also corresponds to the rapid tempo of morphological evolution ^55^ that characterizes the ceratioid backbone, and confirms the evolutionary history of deep-sea anglerfishes features classic signatures of adaptive radiation ^9^.

### Ecological opportunity drove the adaptive radiation of ceratioid anglerfishes

What could have enabled deep-sea anglerfishes to make a major transition into water column habitats and diversify during a period of climate turmoil? To answer this question, we investigated the evolution of the unconventional physiology and behavior of anglerfish reproduction. The sexual parasitism of ceratioid anglerfishes is theorized to increase the chances of successful reproduction after finding a mate in the deep open ocean ^13,22–24,56^. Yet, the association between reproductive mode and lineage diversification in anglerfishes has not been tested. Our analyses reveal significant associations between net lineage diversification and both reproductive mode and habitat in ceratioid anglerfishes (χ^2^ p < 2.2e-16; Table S1; Figure 2A, B).

**Figure 2.**
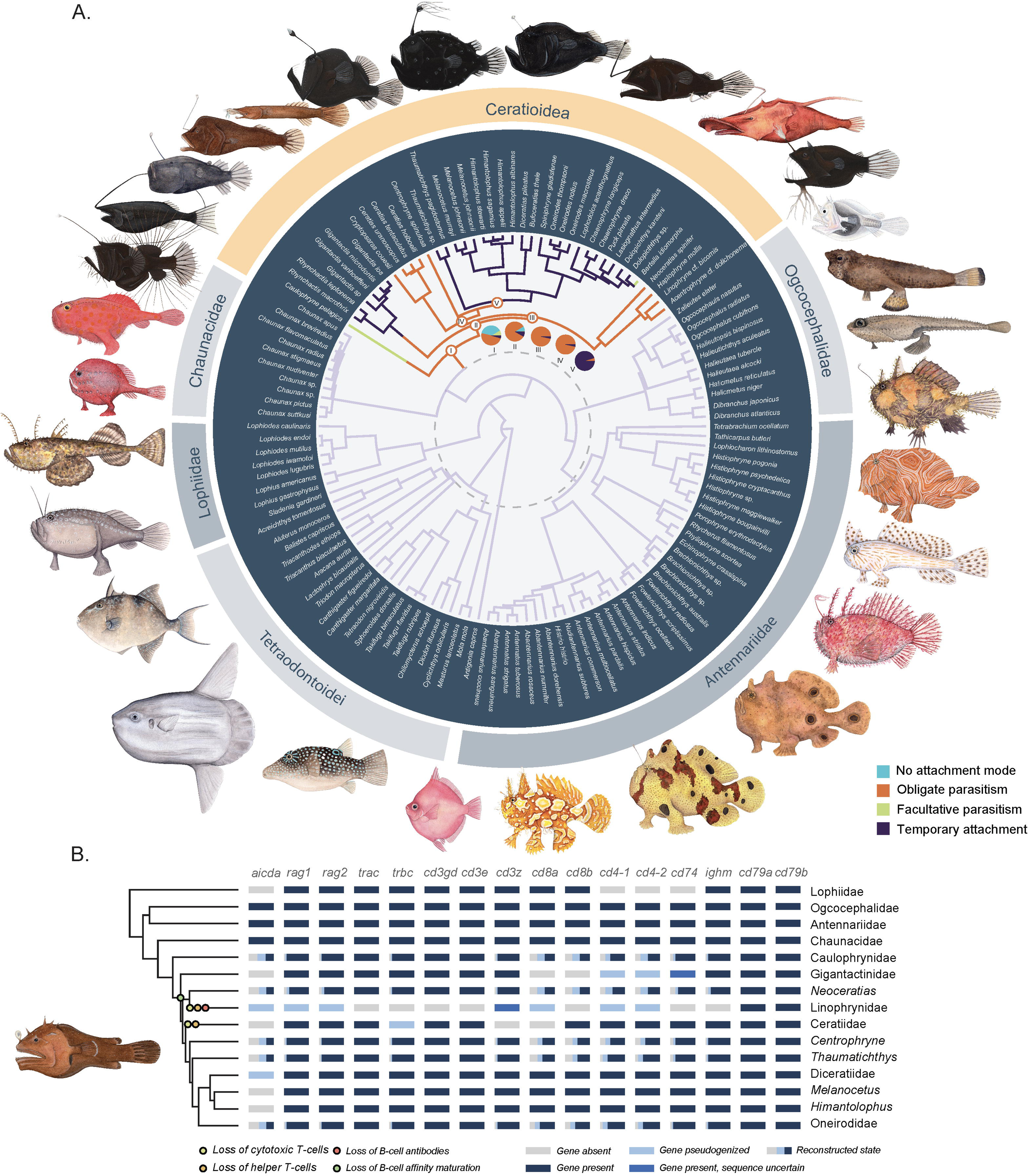
Ecological and reproductive innovations drove the diversification of anglerfishes. Plots show posterior probability density distributions and transition matrices of net diversification rates in anglerfishes with distinctive ecologies and reproductive modes. Panels (A) and (B) show binary state-dependent diversification rate (BiSSE) runs and panels (C) and (D) show multistate-dependent diversification rate (MuSSE) runs for three states. Note that bathypelagic anglerfishes have higher diversification rates than benthic species (B, D), especially compared to deep-sea benthic forms (D). This is driven mainly by diversification associated with the evolution of temporary attachment (A, C).

In contrast, trait-dependent diversification rate analyses using the Binary and Multiple State Speciation and Extinction models ^57^ reveal that clades possessing true sexual parasitism exhibit a similar diversification rate than non-ceratioid anglerfishes (Figure 2C). Temporarily attaching anglerfish clades show the highest lineage diversification rates (Figure 2A, C). Binary trait-based diversification rate analysis show a clear difference in the diversification rates of benthic and bathypelagic clades (Figure 2B). Although multistate analysis of habitat-based diversification suggests that shallow water, predominantly near-shore and reef-dwelling species and the bathypelagic ceratioids have similar diversification rates (Figure 2D), bathypelagic anglerfishes have higher diversification rates than benthic deep-water species, even though benthic deep water species constitute a large proportion of the species diversity of batfishes, goosefishes, and sea toads. These results suggest the secondary invasion of the open waters of the deep ocean–the bathypelagic zone–drove the radiation of ceratioid anglerfishes (Figure 2).

### The evolutionary context of reproductive innovation

Whether obligate parasites resolve as a clade among ceratioid anglerfishes has remained unresolved over half a century of investigations ^13,23,27,27–29,31,40,42^, and recent phylogenomic analyses have failed to reach a consensus ^13,27,28^. Our species tree and maximum likelihood phylogeny fail to recover a single origin for obligate parasitism, contrasting with recent analyses^28^. Ancestral state reconstructions of reproductive mode on the time-calibrated phylogenies of anglerfishes and tetraodontoids support a complex evolutionary history of sexual parasitism. Sexual parasitism was inferred as the state for the common ancestor of ceratioids (Figure 3A). Temporary attachment is reconstructed with two origins: once in Gigantactinidae and at the MRCA of the clade that includes *Thaumatichthys*, *Himantolophus*, Oneirodidae, Diceratiidae, and *Melanocetus* (Figure 3A). All nodes in the anomaly zone are reconstructed as ancestrally sexual parasites with varying degrees of support (Figure 1, Figure 3A). The lack of phylogenetic resolution may be the result of incomplete lineage sorting driven by a history of short intervals between divergence events ^38,46,47^, suggesting that periods of rapid lineage origination were paired with innovation in reproductive mode in deep-sea anglerfishes.

**Figure 3.**
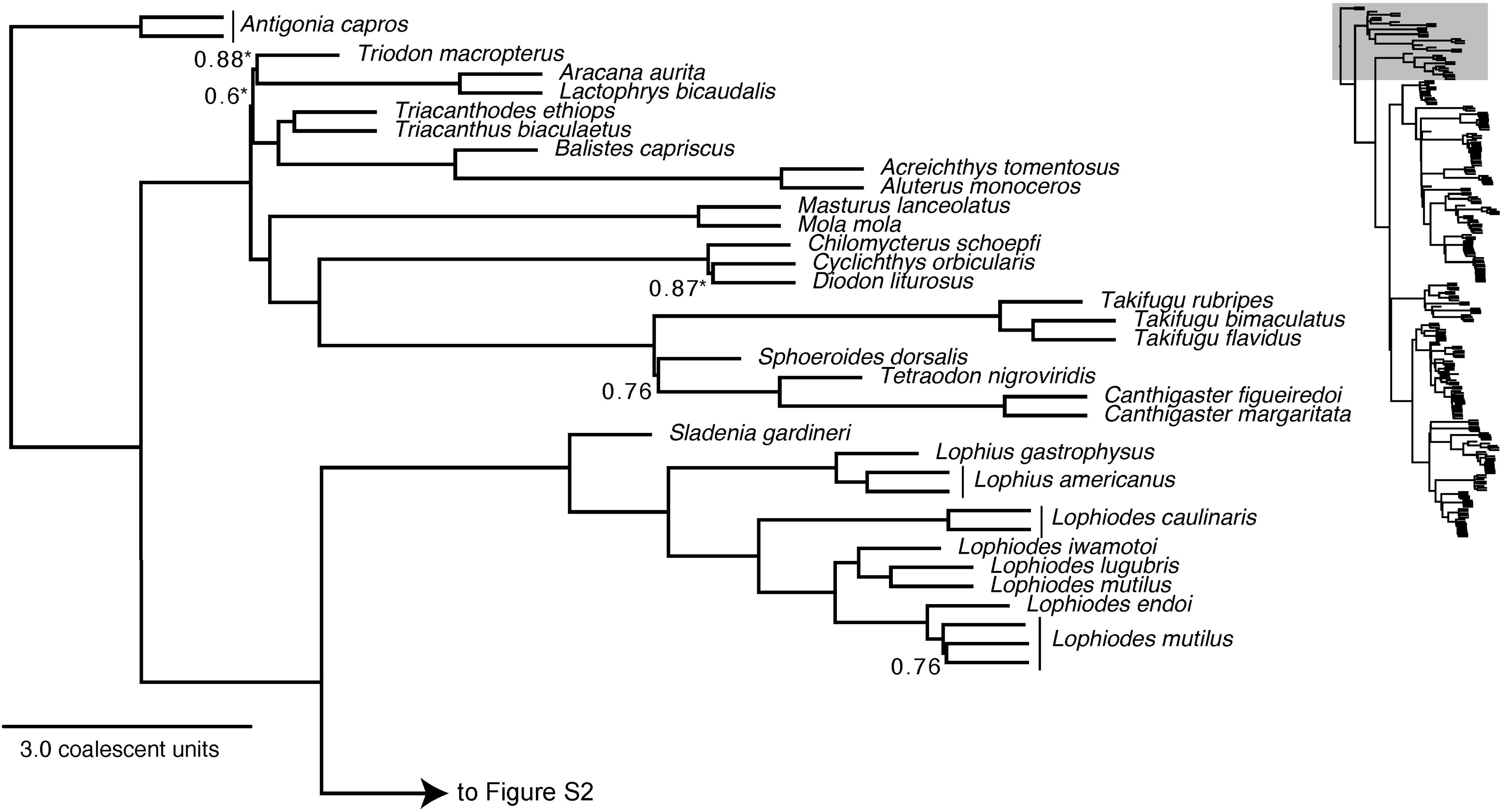
The evolutionary context of anglerfish immunogenomic degradation. Bayesian timetree in (A) shows an ancestral state reconstruction of reproductive mode in anglerfishes and their sister clade, the Tetraodontoidei, based on simulated stochastic character mapping. The dotted line indicates the Cretaceous-Paleogene boundary, and pie charts indicate ancestral state probabilities at key nodes where ancestral states are most uncertain. Light blue indicates no modified reproductive mode, yellow indicates facultative sexual parasitism, orange indicates obligate sexual parasitism, and purple indicates temporary attachment. Fish illustrations by Julie Johnson. Time calibrated phylogeny and table in (B) shows the gene losses and pseudogenizations of key immune genes in anglerfishes based on genomic sequences and simulated stochastic character mapping. Note how different gene losses and pseudogenization events occur in nearly every lineage that displays obligate or facultative sexual parasitism.

Recent work has shown that both temporarily attaching and sexually parasitic ceratioid anglerfishes possess degenerate genomic bases of adaptive immunity ^13^. However, different hypotheses of deep sea anglerfish phylogeny ^13,27–29,40^ imply different modes of immunogenomic change. We compiled data on the presence of selected genes relevant to adaptive immunity in anglerfishes from the literature ^13,25^, GenBank searches, and BLAST runs of published anglerfish genomes against identified sequences of interest in lophioids ^13,58^, and then used multiple methods of ancestral state reconstruction to produce gene presence and absence likelihoods for clades lacking sequence data (Figure 3B). Ancestral state reconstructions show that cellular components of adaptive immunity convergently degenerated in clades with parasitic reproductive modes (Figure 3B; Figure S12) due to parallel changes in their immunogenomics (Figure 3B), rather than via the stepwise loss or pseudogenization of genes along the backbone of the deep sea anglerfish clade. Further, non-ceratioid lineages such as batfishes and goosefishes demonstrate complex losses and pseudogenizations of key immune genes of non-ceratioid anglerfishes (Figure 3B). Whereas ogcocephalids, chaunacids, and antennariids show no losses to their immunogenomic loci, lophiids have lost virtually all of the genetic basis of the major histocompatibility complex II ^25^. These observations, particularly when considered in the context of the various episodes of immunogenomic degeneration in teleost fishes ^21,59^, show that modifications and losses of key genes involved in adaptive immunity predated the origins of deep-sea anglerfishes. Similarly, ancestral state reconstructions for sexual body size dimorphism based on recent observations of diminutive males in frogfishes and the deep sea lophiid *Sladenia gardineri* ^60^ suggest that significant body size differences between male and female anglerfishes originated prior to the origin of deep-sea ceratioids (Figure S13).

## Discussion

Anglerfishes comprise the most species-rich radiation of vertebrates in the deep sea ^13,24,27,29,40,42^, an evolutionary feat that may have been aided by peculiar reproductive modes that increased the chances of successfully mating in the largest of Earth’s ecosystems ^23,24,61^. Here, we provide a phylogenomic reconstruction of the evolutionary history of anglerfishes. By examining the evolutionary history of genes encoding critical aspects of the adaptive immune response and the evolution of sexual dimorphism in all major anglerfish lineages using a time-calibrated phylogenomic tree, we determined that degeneration of the genomic basis of adaptive immunity and extreme size differences between male and female anglerfishes coincident with the origin of ceratioids and the evolution of sexual parasitism (Figure 3b)^25^. Our results suggest that the high synergy between these traits may promote the persistence of unique pathways to degeneration (Figure 3b) that different clades of obligate parasite ceratioids possess ^13^. Collectively, these results suggest the development of new modes of reproduction in deep-sea anglerfishes was a synergistic process involving the loss of core immunogenetic functionality and extreme sexual size dimorphism (Figure 3). However, it was the confluence with a major ecological transition into the open ocean during a period of intense global ecological stress that facilitated higher rates of diversification in bathypelagic lineages.

Our results implicate the degradation of the genetic basis of adaptive immunity and extreme sexual dimorphism as mechanisms driving the ecological transition of ceratioid anglerfishes into deep-sea open water habitats. We reveal that AICDA was lost prior to the MRCA of Ceratioidea, potentially compromising the ability to generate high affinity antibodies or switch between antibody isotopes (IgT, IgM, etc; ^62^ Figure 3B). This timing is coincident with a dramatic shift towards extreme size dimorphism between male and female ceratoid anglerfishes (Figure S13). Diminutive males of non-ceratioid anglerfishes are known to closely follow the larger females ^60^, suggesting behaviors reminiscent of temporary attachment are much more widespread within Lophioidei. In lineages such as the obligate sexual parasites Linophrynidae, the lack of a functional antibody response likely set the stage for a suite of losses otherwise critical for the adaptive immune response including V(D)J recombination mediated by RAGI and RAGII, MHCII antigen display with CD74, and T-cell functionality that now enables fusion of male anglerfish to females. Together with ancestral state reconstructions (Figure S13), these observations indicate sexual parasitism was assembled from a suite of genomic, morphological, and behavioral characteristics with evolutionary origins stretching back into the Mesozoic (Figure 1, Figure 3).

There is quantifiably unstable phylogenomic backbone coincident with the origin and initial diversification of ceratioid anglerfishes (Figure 1), which radiated into open water habitats from benthic fin-walking ancestors (Figure 1, Figure 2B). However, such uncertainty is expected if anglerfishes radiated at the confluence of the Paleogene-Eocene Thermal Maximum. The extensive gene tree discordance and short branch lengths across the ceratioid phylogenomic backbone are indicative of ancestral polymorphism resulting from an ancestral large effective population size and relatively short intervals between speciation events ^38,46,47^ (Figure 1; Figure 2B, D). Rather than resulting from poor genetic sampling or methodological artifacts, the anomaly zone at the base of ceratioids appears to reflect an evolutionary history of relatively rapid lineage diversification. Similarly discordant nodes have been found in clades like birds that underwent adaptive radiation following major extinction events ^63–66^. The time-calibrated phylogeny generated in this study shows that nearly all ceratioid lineages classified as taxonomic families appeared during the Paleogene-Eocene Thermal Maximum (Figure 1), an interval marked by major marine turnovers and extinction events, including lineages of the Tetraodontoidei, the sister clade to anglerfishes ^49^.

Nearly all examples of evolutionary radiations highlight traits that are held responsible for directly promoting specialization in new ecological niches. Classic examples include the fused pharyngeal jaws of cichlid fishes ^67–72^ and the beaks of Darwin’s Finches ^73–75^. Our results suggest that extreme sexual size dimorphism and loss of the adaptive immune response form the synnovation that gave rise to sexual parasitism, a trait that facilitated access to a previously inaccessible ecosystem by ensuring reproductive success ^7^. However, examination of trait-dependent diversification rates suggests that ceratioid diversification ^55^ was primarily driven by ecological opportunities available in the bathypelagic zone (Figure 2) during the Paleocene-Eocene Thermal Maximum (Figure 1) rather than by their bizarre reproductive strategies. In this way, the sexual parasitism of ceratioid anglerfishes provides an apt example of key innovation as the term was originally conceived ^7,76,77^ and shows how these traits need not be solely responsible for the evolutionary diversification of a lineage. A complex evolutionary cascade of many different traits, including sexual dimorphism, degeneration of many key pathways of the adaptive immune system, and the evolution of a body plan suited for a pelagic lifestyle, allowed anglerfishes to diversify in the bathypelagic zone as oceanic ecosystems faced extreme global warming.

## Methods

### Taxon Sampling and Sequencing Methods

In order to effectively sample lophioid diversity, we sequenced and aligned 1,314 UCE loci for 230 specimens of anglerfishes and 22 outgroups (2 individuals of *Antigonia capros* and 20 tetraodontoids representing all major clades) ^28,78^.

*Lophichthys boschmai*, which traditionally comprises the monotypic Lophichthyidae, was not sampled. However, this species is nested within Antennariidae (*sensu* ^28^) in morphological analyses in which it is included ^79,80^. As such, our sampling can be said to comprise 100% of lophioid families, ∼70% of genera, and ∼30% of the species in the clade. The UCE data for 157 specimens come from previous studies ^27,28^. New sequence data was generated following protocols outlined in previous phylogenomic analyses of acanthomorph teleosts ^28,81^. We isolated DNA from muscle and fin tissue using Qiagen DNeasy Blood and Tissue kits, quantified 1µl for each extraction using a Qubit fluorometer (Life Technologies), confirmed successful isolation of high molecular weight DNA using gel electrophoresis, and sheared ∼500 ng of DNA for each sample using a QSonic Q800R3 sonicator to produce sequence fragments ranging in length between 300 and 600 nt. Using Kapa HyperPrep kits (Kapa Biosystems) and Illumina TruSeq iTru5 and iTru7 adapters ^82^, we indexed genomic libraries. During all purification steps, we conducted dual-step SPRI bead cleanups using 80% EtOH washes. This protocol included a 0.8X clean-up after adapters were ligated. We used 25 μl KAPA HiFi HotStart ReadyMix (Kapa Biosystems), 5 μl of 5uM Illumina TruSeq iTru5 and iTru7 dual-indexed primers ^82^, and 5 μl ddH_2_O in a thermocycler with the following settings to amplify DNA: 45 s at 98° C, 13 cycles for 15 s at 98° C, 30 seconds at 60° C, and 60s and a subsequent final extension at 72° C. PCR products were purified using 1.0X SPRI beads. Libraries were then rehydrated in 25 μl of 10 mM Tris-HCL and quantified using a Qubit fluorometer. We pooled 80 ng of each sample into groups of nine, dried in a vacuufuge, and rehydrated using 4.9 μl 10 mM Tris-HCL. Using a published bait set for 1314 UCE loci ^28,78^, we used the following targeted enrichment protocol: pooled libraries were hybridized with 100 ng of Arbor Sciences myBaits, 500 ng custom blocking oligoes, and 1% sodium dodecyl sulfate at 65° C for 24 hours. We followed with 2 bead clean ups on all pools, removed remaining wash buffer, dried samples on the magnetic rack, and then rehydrated them with 30 μl ddH_2_O. After combining bead-bound enriched libraries with 25 μl Kapa Biosystems Hifi HotStart ReadyMix polymerase, with 5 μl Illumina TrueSeq primer mix and 5 μl ddH_2_O each, we ran PCR on the samples with 16 amplification cycles and the same time and temperature settings as our amplification protocol. After reaction products were purified using single bead clean-ups, we rehydrated pools using 33 μl ddH_2_O, quantified them, diluted them to 2.5 ng μl^-1^, used an Agilent Technologies Bioanalyzer to confirm their size distribution, quantified each pool using a Kapa Biosystems qPCR kit, and adjusted sample concentrations to 10 nM using Bioanalyzer mean fragment size estimates and our qPCR results. We then sequenced the libraries using 150 bp paired-end sequencing on Illumina NovaSeq platforms after creating an equimolar pool of enriched libraries.

### UCE Sequence Alignment, Sanitation, and Preprocessing

We used the phyluce 1.7.1 pipeline^32^ to process raw reads, assemble sequences *de novo*, and construct UCE sequence alignments. Phyluce was also used to check for and remove potential paralogs. This produced 975 UCE alignments. We conducted several additional steps to identify and filter paralogous and chimeric sequences from our dataset. First, we used CIAlign ^33^ to flag and visualize chimeric or incorrectly included UCE sequences produced by phyluce. Next, we used custom scripts in the R package TOAST ^36^ to conduct a secondary screen for hidden paralogous genes and anomalously divergent sequences by checking for reciprocal monophyly of Lophioidei and the two outgroup clades (Tetraodontoidei and *Antigonia capros*). Finally, we tested for violations of common sequence substitution model assumptions, including stationarity, homogeneity, and reversibility, using scripts ^83^ implemented in IQ-TREE 2.2.03 ^34,35^. The final set of UCEs comprised 608 aligned loci for 222 individual specimens that comprise 124 species.

### Maximum Likelihood Phylogenetic Analyses

All maximum likelihood phylogenetic analyses were conducted on the Yale High-Performance Computing McCleary and Farnam Clusters using IQ-TREE v. 2.2.0 ^34^. We conducted three separate maximum likelihood phylogenetic analyses that constructed final species trees using different methodologies. First, we inferred gene trees for individual UCE sequences using 1000 ultrafast bootstrap supports, 1000 replicates of the Shimodaira–Hasegawa approximate likelihood ratio (SH-r) test, and models of nucleotide evolution for individual UCEs found using ModelFinder Plus. These were summarized into a species tree using ASTRAL III v. 5.7.8 ^37^. Next, we partitioned the UCE alignments into 41 sets using PartitionFinder2 ^84^ and inferred a maximum likelihood topology using 1000 ultrafast bootstraps. We inferred a maximum likelihood tree for the whole concatenated alignment using standard bootstraps and a model of nucleotide evolution inferred using ModelFinder Plus.

### Concordance Factors and Anomaly Zone Detection

The magnitude of gene tree-species tree discordance was assessed by calculating gene (gCF) and site (sCF) concordance factors using IQ-TREE 2 ^34^. Values of gCFs measure the number of decisive gene trees consistent with particular branches in the partitioned tree topology, and sCFs measure the number of decisive alignment sites supporting branches in the partitioned tree. We calculated sCFs using 100 subsampled quartets from our alignment and plotted gCF, sCF, bootstrap, and branch length values against each other to test for associations using R base code and tidyverse scripts (http://www.robertlanfear.com/blog/files/concordance_factors.html). Examination of these plots revealed two regions of phylogenetic instability, one corresponding to most of the backbone divergences in *Ceratioidea* and another corresponding to genus-level divergences in *Antennariidae*. To further interrogate our data and locate discordant regions of our generated phylogeny, we used custom scripts on the ASTRAL species tree to test for the presence of anomaly zones at particular regions of the tree. Anomaly zones (AZs) can be detected by checking whether branch lengths scaled to coalescent units fall within theoretical boundaries for AZs ^38,46,47^; we transformed branch lengths to coalescent units using quartet CFs before testing whether these branches were AZs using custom python code ^46,47^.

### Divergence Time Estimation

We used a node-dating approach implemented in BEAST 2.6.6 ^48,85^ to time-calibrate the phylogeny of lophioids using input xml files of the 15 largest partitions of UCE loci that we delimited using IQ-TREE 2. Input files were constructed using BEAUTi.^48,85^ We used the General Time Reversible (GTR) model with gamma distributed among site rate variation model of nucleotide substitution, a relaxed log-normal molecular clock model, and the Fossilized Birth-Death (FBD) model of divergence time estimation as implemented in BEAST 2^86^. For the FBD model, we specified rho as 0.25, which is the sampling fraction of all 408 recognized species of *Lophioidei* (following Eschmeyer’s Catalogue of Fishes: https://www.calacademy.org/scientists/projects/eschmeyers-catalog-of-fishes) included in our dataset. We set the origin to 83.6 Ma as a uniform prior, which is the age of the oldest occurrences of fossils from the major crown percomorph clades (e.g., †*Gasterorhamphosus zuppichinii* ^28,78,87^, †*Nardoichthys francisci* ^88^), with bounds of 145.0 Ma (Jurassic-Cretaceous boundary, marking the start of the fossil record and estimated ancestry of crown Acanthomorpha ^28,31,41,78,89,90^) and 56.0 Ma (the oldest crown tetraodontoid fossils, marking the minimum age of the tetraodontoid-lophioid split) ^30,54,91^. The diversification rate parameter was set to 1.47, which is the number of lophioid species included in our dataset divided by the median age of the root origin prior, with bounds of 0 and 100.0. We used eleven fossil calibrations to node-date the tree, each of which are discussed in the Fossil Calibration Justifications section of the Supplementary Text. We fixed the tree topology to the maximum likelihood phylogeny from the partitioned IQTREE analysis. Each partition was analyzed in BEAST for 200 million generations with a 10 million generation pre-burnin. Convergence of the posteriors was assessed using Tracer v. 1.7.1 ^92^ and converged runs were combined using LogCombiner 2.6.4 (subsampling every million generations) ^85^ and annotated to the target IQTREE topology using TreeAnnotator 2.6.6 ^85^ using median node heights.

### Ancestral State Reconstructions

We implemented two methods of discrete trait analysis to reconstruct the ancestral states in the evolutionary history of locomotory and reproductive modes in anglerfishes using the R packages ape ^93^ and phytools ^94^. Reproductive modes were classified as temporary attachment, facultative parasitism (both temporary attachment and permanent fusion), obligate parasitism (permanent fusion only), or no reproductive parasitism. Data on locomotory and reproductive modes were taken from four primary studies ^22–24,61^. We do note that at least one study has argued that all ceratioids might be parasitic ^61^ and that temporary attachers are just facultative species for which parasitism has not yet been observed. However, the genomic evidence and our ancestral state reconstructions of gene presence (see below) suggests that at the very least, if this is true, the majority of anglerfishes would be facultatively parasitic without a degenerated genetic basis of major features of adaptive immunity. As such, we keep the standard classification of anglerfishes as either temporary, obligate, or facultative parasites.For data on sexual dimorphism, we assembled information on size differences in females and males from the literature ^22–24,56,60,61,80^. In each case, we first used the fastAnc function to produce a rapid maximum likelihood estimate of ancestral states at nodes in the time-calibrated phylogeny. The make.simmap function was used to fit a continuous-time reversible Markov model on the evolution of reproductive mode and simulate stochastic character evolutionary histories over 1000 simulations using the Markov model and our inputted species trait data. The results were summarized on a single tree.

### Chaunacid Mitochondrial Gene Tree and Genetic Distances

For most of the major lineages of anglerfishes, we are confident that our species sampling captures the common ancestor node. The exception is the *Chaunacidae* (sea toads). The UCE phylogeny includes one of two genera and 10 of 25 recognized species of *Chaunax*, we cannot be certain that the common ancestor of crown lineage of *Chaunacidae* is included in the tree. The time-calibrated UCE phylogeny indicates that crown *Chaunax* diversified recently relative to all other lophioid genera, so we suspected this genus is taxonomically over split. These observations posed challenges to the use of a speciation-based branching model clade sampling fractions into relaxed clock and diversification analyses. We investigated the delimitation of sampled species of *Chaunax* by sampling sequences of the mtDNA gene *COI* for 11 of the 25 recognized species of *Chaunax* and one of four species of *Chaunacops*. The *COI* DNA sequence alignment was analyzed using IQ-TREE with ultrafast bootstrap supports generated over 1000 replicates, 1000 replicates for the SH-r test, and model specification in ModelFinder Plus. The resulting phylogeny was rooted on *Chaunacops coloratus.* Finally, we imported the *COI* DNA sequence alignment into R and calculated pairwise genetic distances across *Chaunacidae* using the R package ape.

### Trait-Based Species Richness and Diversification Rates

We conducted several tests for associations between species richness and reproductive and locomotor traits in lophioids. First, we used chi-square tests implemented in R to investigate associations between species richness, reproductive mode, locomotory mode, and habitat in anglerfishes. Species diversity reflected the number of recognized species reported in Eschmeyer’s Catalogue of Fishes and categorized for each trait state by reviewing the literature ^13,29,39,61^. From these data, we constructed a four-by-three table listing the number of species displaying each of four reproductive modes: no sexually parasitic reproductive mode (1), sexually parasitic reproductive mode (2), and temporary attachment (3); and three different habitat modes based on data from FishBase.se: (1) shallow water benthic (<200 m), (2) deep water benthic (>200 m), and (3) bathypelagic. We used the Rstats function chisq.test to run χ2 and examined expected, observed, and residual values. Next, we conducted trait-dependent diversification rate analyses using the R package diversitree^57^ using MuSSE models for the reproductive and locomotory mode character sets. We tested for differences in diversification rates associated with locomotory modes by running a BiSSE (binary state dependent diversification rate analysis) on locomotory mode characterized as either demersal or open-water-swimming. For the MuSSE analyses of the whole sample, reproductive mode was categorized as either (1) no sexual parasitism, (2) true sexual parasitism, or (3) temporary attachment; for MuSSE analyses on reproductive mode among ceratioids and locomotory mode, trait categories were the same as those in the χ2 tests. In each SSE analysis, sampling fractions for each trait were set based on the number of species represented in the phylogeny relative to the clade species diversity reported in Eschmeyer’s Catalogue of Fishes. In BiSSE, we tested likelihood models where speciation rates were the same or allowed to differ.

In MuSSE, we compared the fits of six different likelihood models variously constraining trait transition rates, speciation rates, and extinction rates to our three-state trait sets using AIC scores. For the best-fit model in each case (the model with the lowest score), we ran an MCMC chain for 10,000 generations with sampling every 100 generations, discarded the first 1000 samples as burn-in, checked for convergence by examining caterpillar plots of log-likelihood scores per generation, and plotted net diversification rate distributions and transition rates among states.

### Evolutionary History of Anglerfish Immunogenetics

Some genetically nonidentical male and female anglerfishes have the ability to fuse because they have degenerated genetic bases for vertebrate adaptive immunity ^13^. To reconstruct the variance in anglerfish immunogenetics and investigate the evolutionary history of these genomic losses, we compiled data on the immunogenetics of key clades *Lophioidei* from recent studies ^25,58^, searches on the NCBI Genbank database, and using the basic local alignment search tool (BLAST) to search published anglerfish genomes for the following core immune genes (Figure 3B) using sequences from *Danio rerio.* These genes were chosen due to their well-known functionality in vertebrates including fishes and their central role in mediating or enabling the adaptive immune response.

This approach allowed us to fill in a table of anglerfish immune gene losses, pseudogenizations, and other modifications. Using these data, we inferred the presence of key features of the adaptive immune system in a subset of anglerfishes and conducted ancestral state reconstructions in the R packages ape ^93^ and phytools ^94^ for the following: cytotoxic T-cells, helper T-cells, B-cell antibodies, and B-cell affinity maturation. We coded three states for each: present, absent, and present but pathway degraded/receptors missing as in a previous study ^13^. We used fastAnc as a first-pass reconstruction followed by make.simmap to fit continuous-time reversible Markov models and simulate stochastic character evolutionary histories over 1000 simulations.

## Funding

This work was supported by the National Science Foundation (IOS1755242 to AD) and the Bingham Oceanographic Fund maintained by the Peabody Museum, Yale University.

## Competing Interests

The authors declare no competing interests.

## Acknowledgements

We thank members of the Near and Dornburg labs for discussions regarding ray-finned fish and anglerfish evolution. For critical tissues we thank T. Sutton, A. Cook, H. Bracken-Grissom, and J. Moore from the DEEPEND project.

## Data Availability

All data, including UCE sequences and all secondary analysis input and output files, will be made publicly available upon formal publication of this article.

---

**Figure S1.**
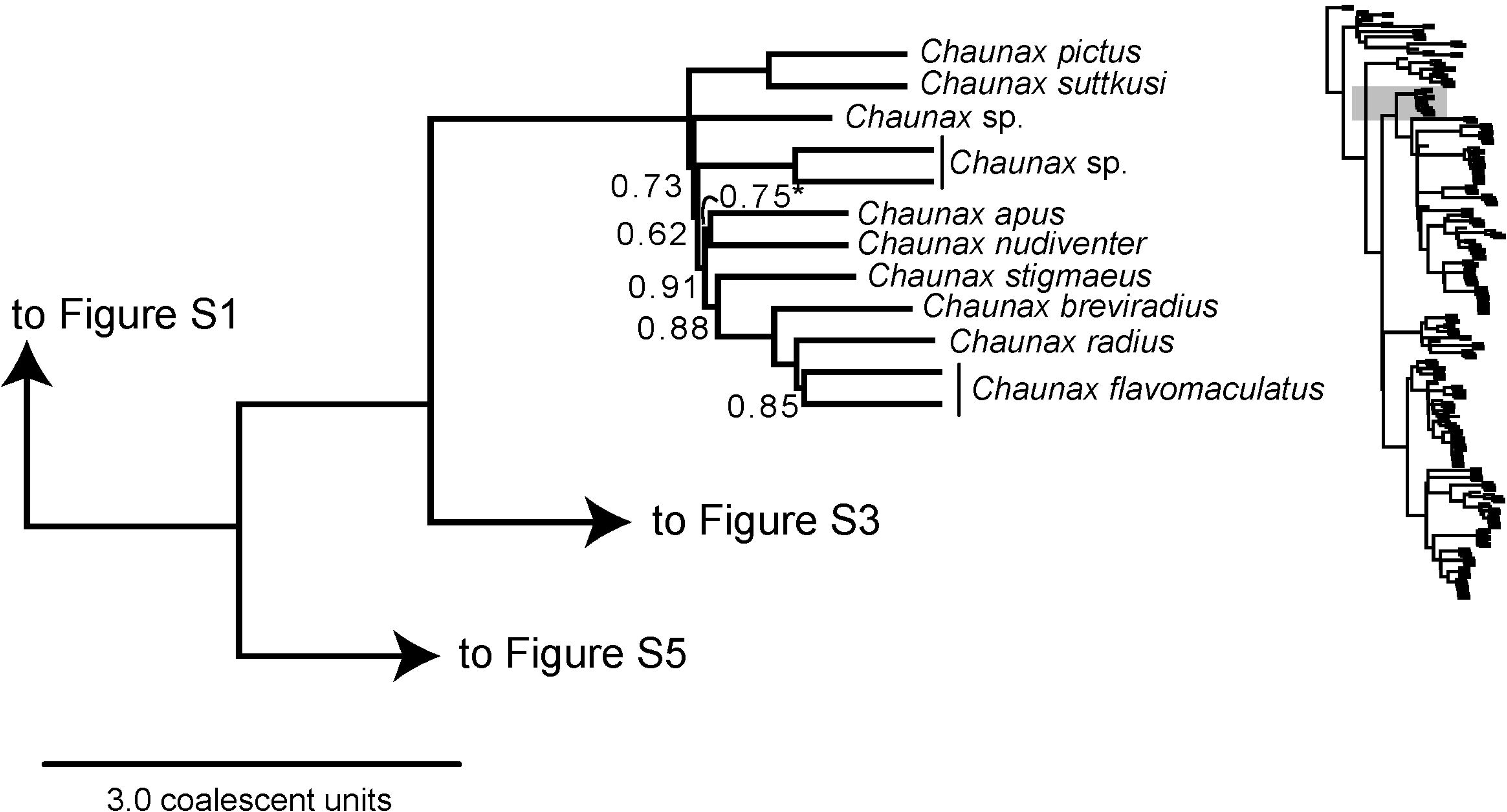
The phylogeny of Anglerfishes I. Inset of ASTRAL III species tree showing relationships of the outgroups (*Antigonia capros* and *Tetraodontoidei*) and *Lophiidae*. Numbers at branches indicate coalescent supports of less than 1.0 Ultimate branches leading to tips have no lengths, but all other branches are scaled to coalescent units.

**Figure S2.**
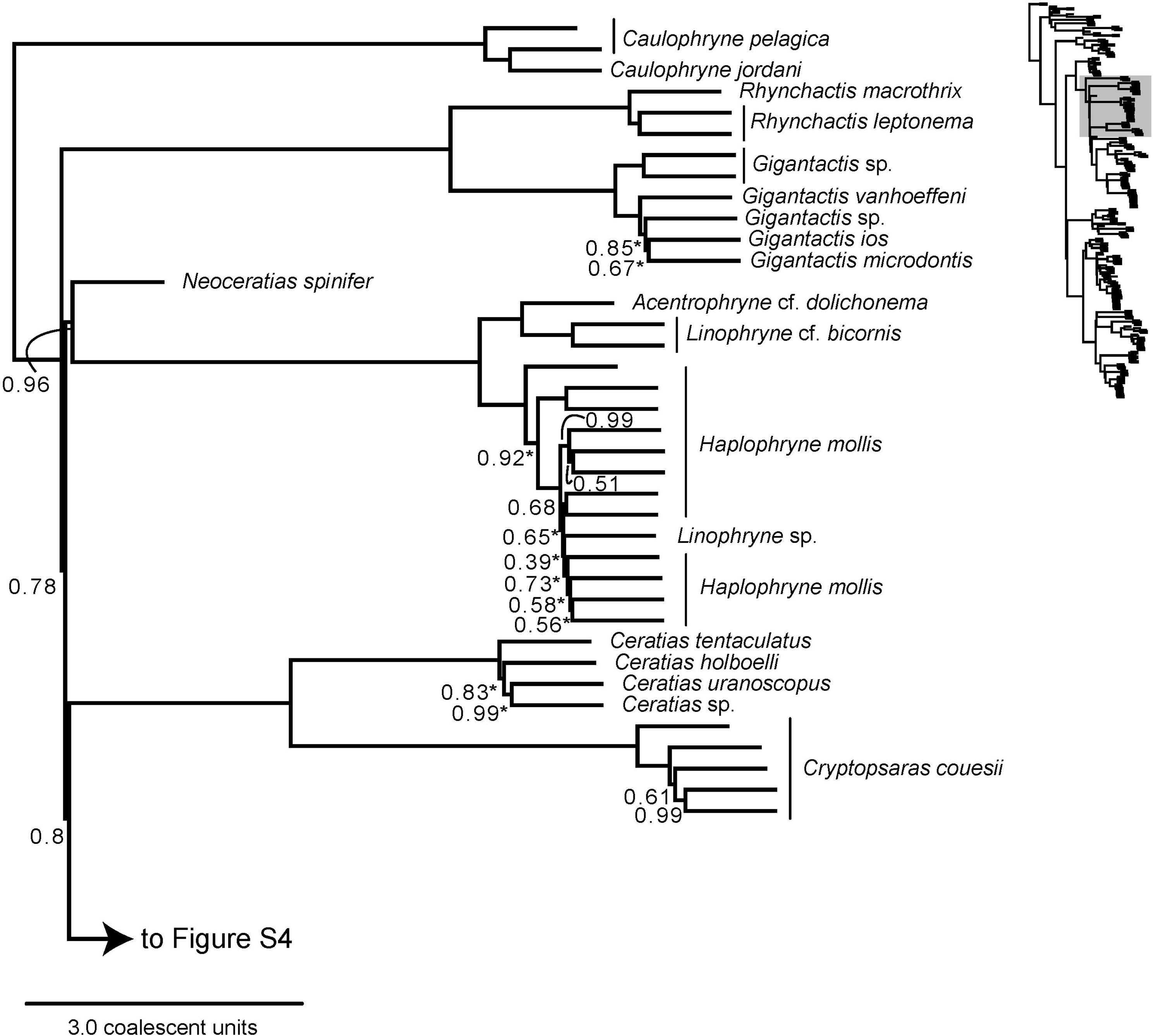
The phylogeny of Anglerfishes II. Inset of ASTRAL III species tree showing relationships of *Chaunaciidae*. Numbers at branches indicate coalescent supports of less than 1.0 Ultimate branches leading to tips have no lengths, but all other branches are scaled to coalescent units.

**Figure S3.**
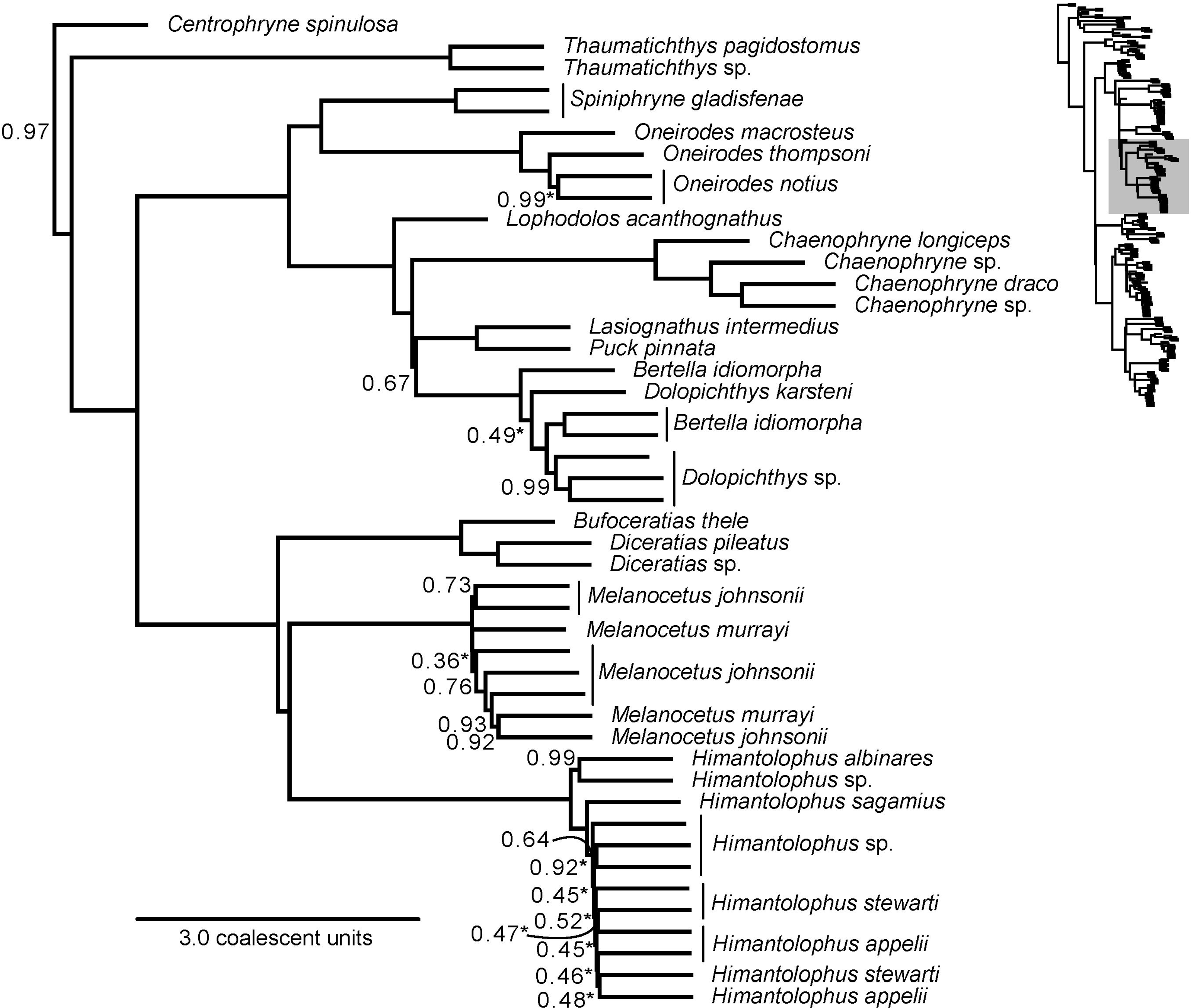
The phylogeny of Anglerfishes III. Inset of ASTRAL III species tree showing relationships of half of *Ceratioidea*. Numbers at branches indicate coalescent supports of less than 1.0 Ultimate branches leading to tips have no lengths, but all other branches are scaled to coalescent units.

**Figure S4.**
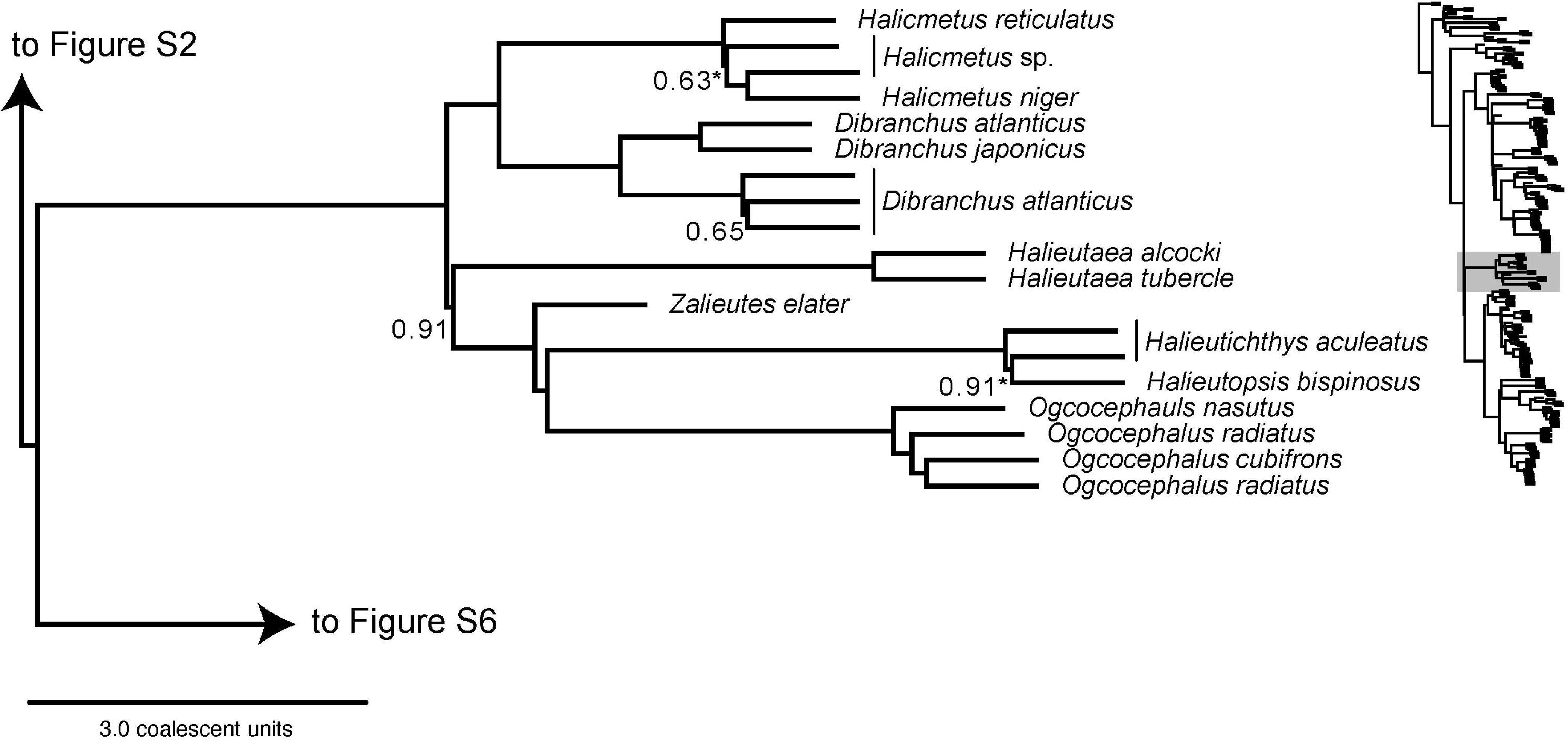
The phylogeny of Anglerfishes IV. Inset of ASTRAL III species tree showing relationships of half of *Ceratioidea*. Numbers at branches indicate coalescent supports of less than 1.0 Ultimate branches leading to tips have no lengths, but all other branches are scaled to coalescent units.

**Figure S5.**
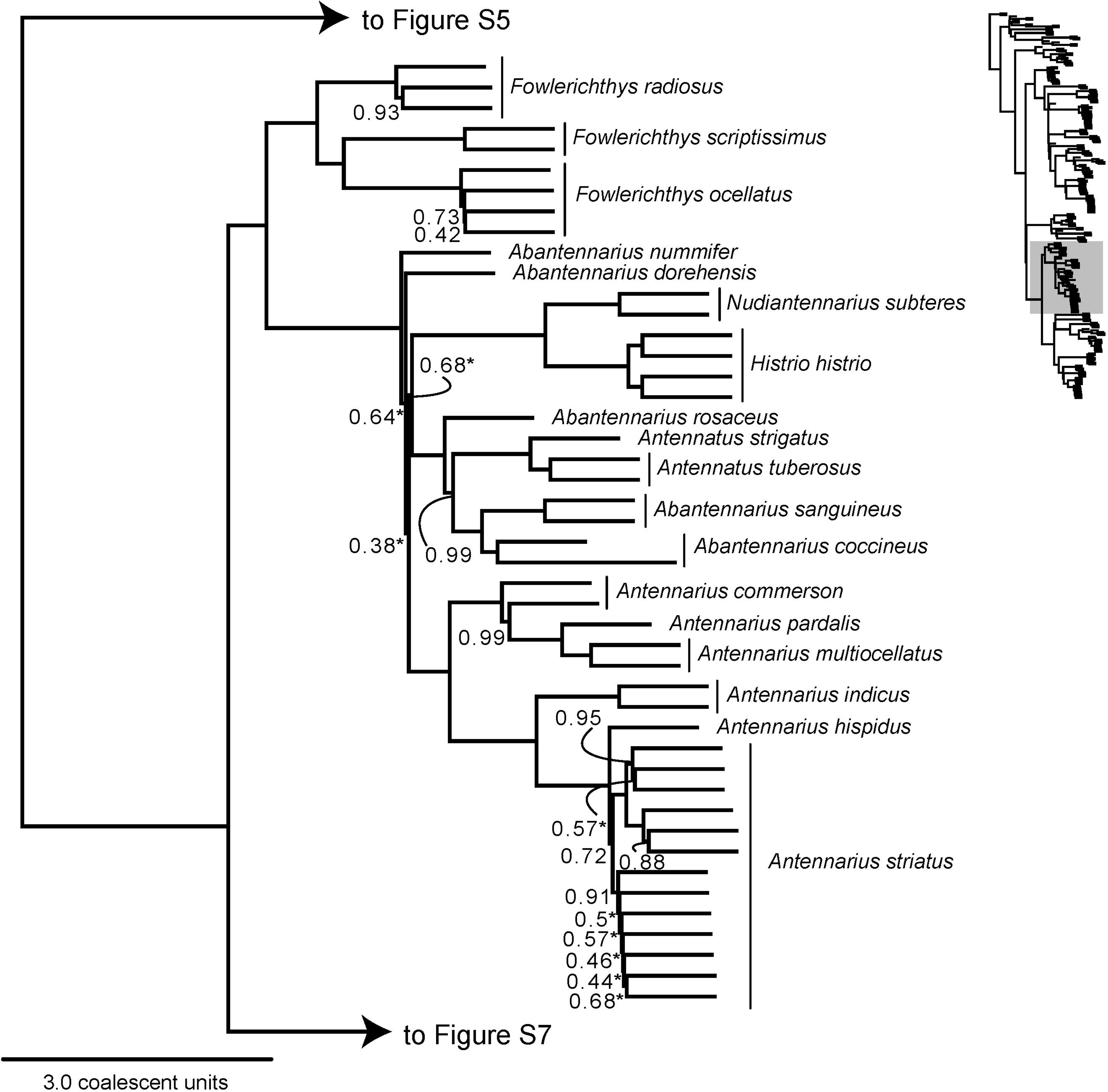
The phylogeny of Anglerfishes V. Inset of ASTRAL III species tree showing relationships of *Ogcocephalidae*. Numbers at branches indicate coalescent supports of less than 1.0 Ultimate branches leading to tips have no lengths, but all other branches are scaled to coalescent units.

**Figure S6.**
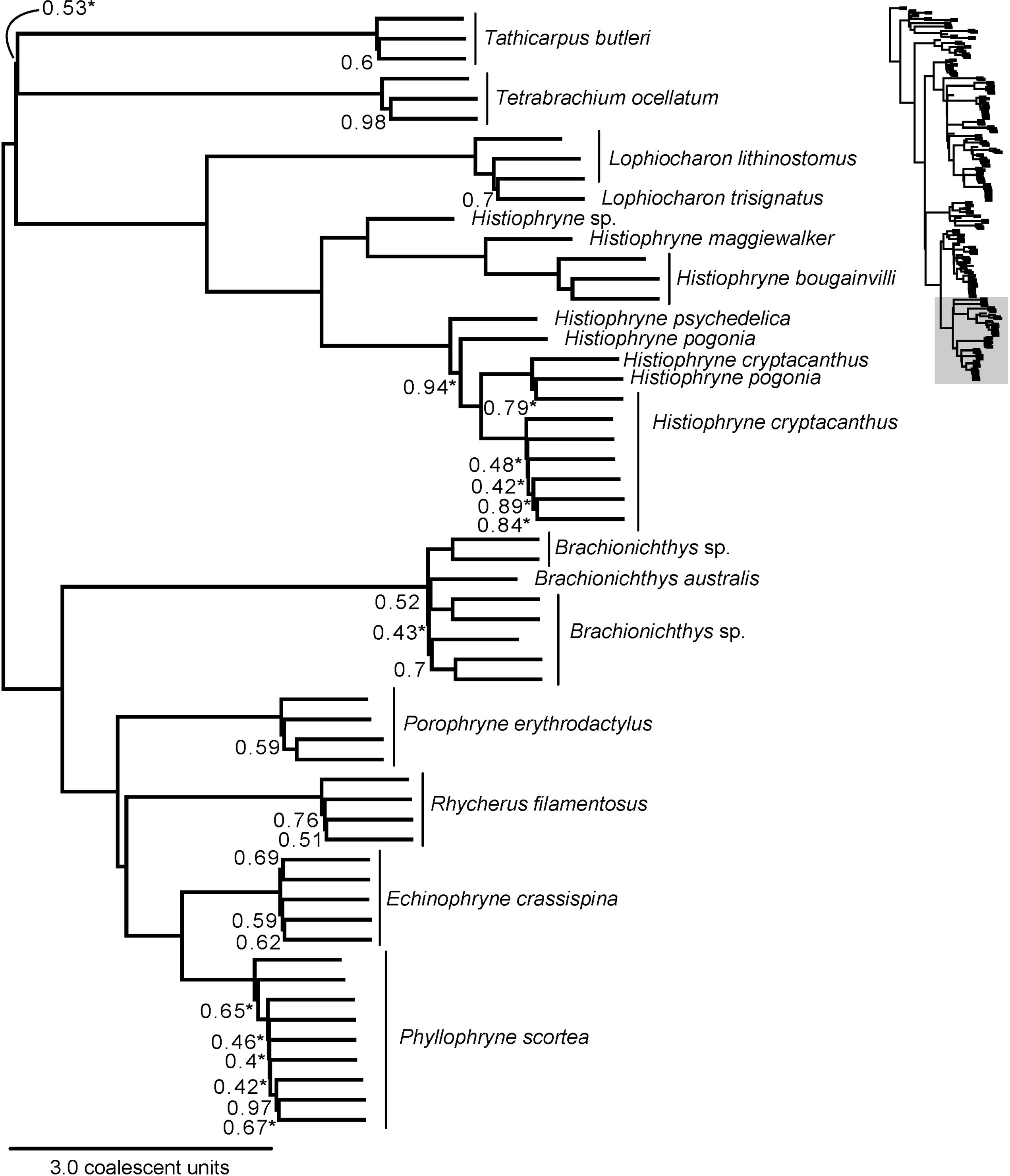
The phylogeny of Anglerfishes VI. Inset of ASTRAL III species tree showing relationships of half of *Antennariidae*. Numbers at branches indicate coalescent supports of less than 1.0 Ultimate branches leading to tips have no lengths, but all other branches are scaled to coalescent units.

**Figure S7.**
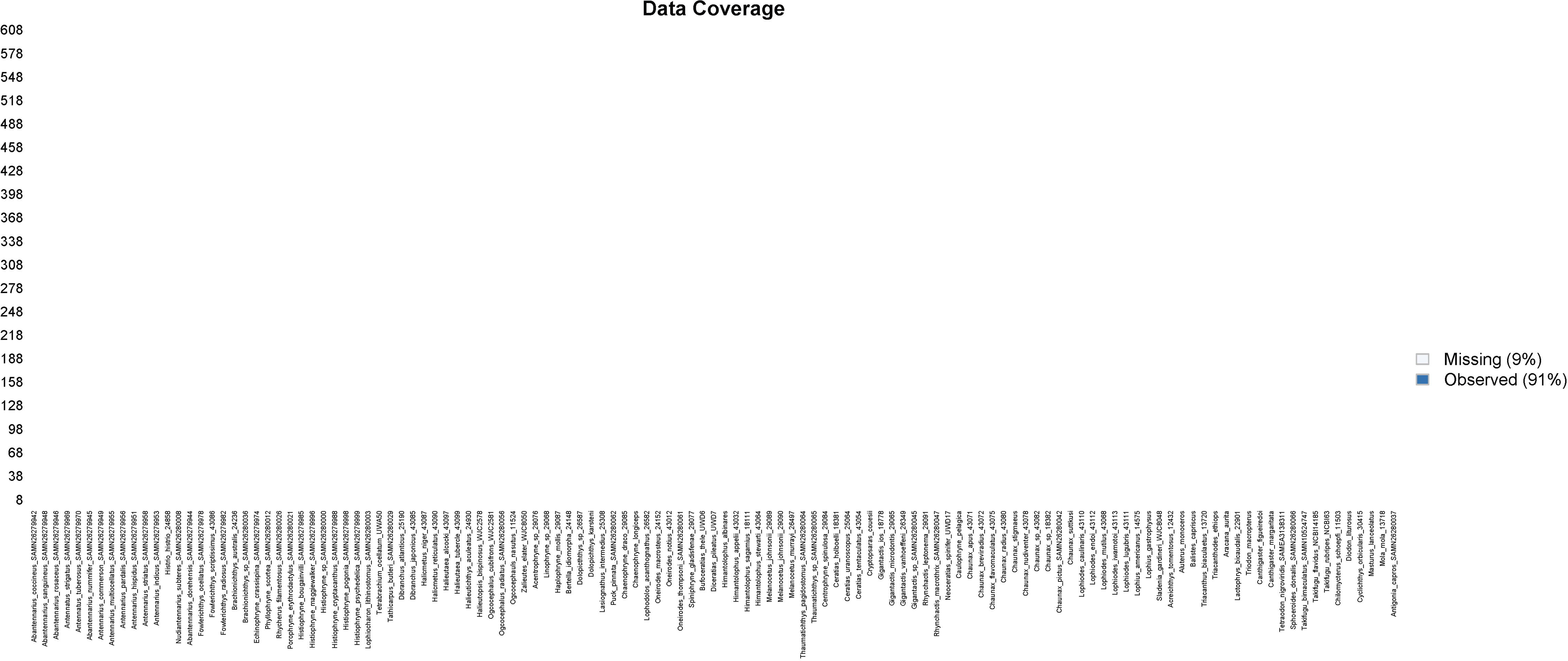
The phylogeny of Anglerfishes VII. Inset of ASTRAL III species tree showing relationships of half of *Antennariidae*. Numbers at branches indicate coalescent supports of less than 1.0 Ultimate branches leading to tips have no lengths, but all other branches are scaled to coalescent units.

**Figure S8.**
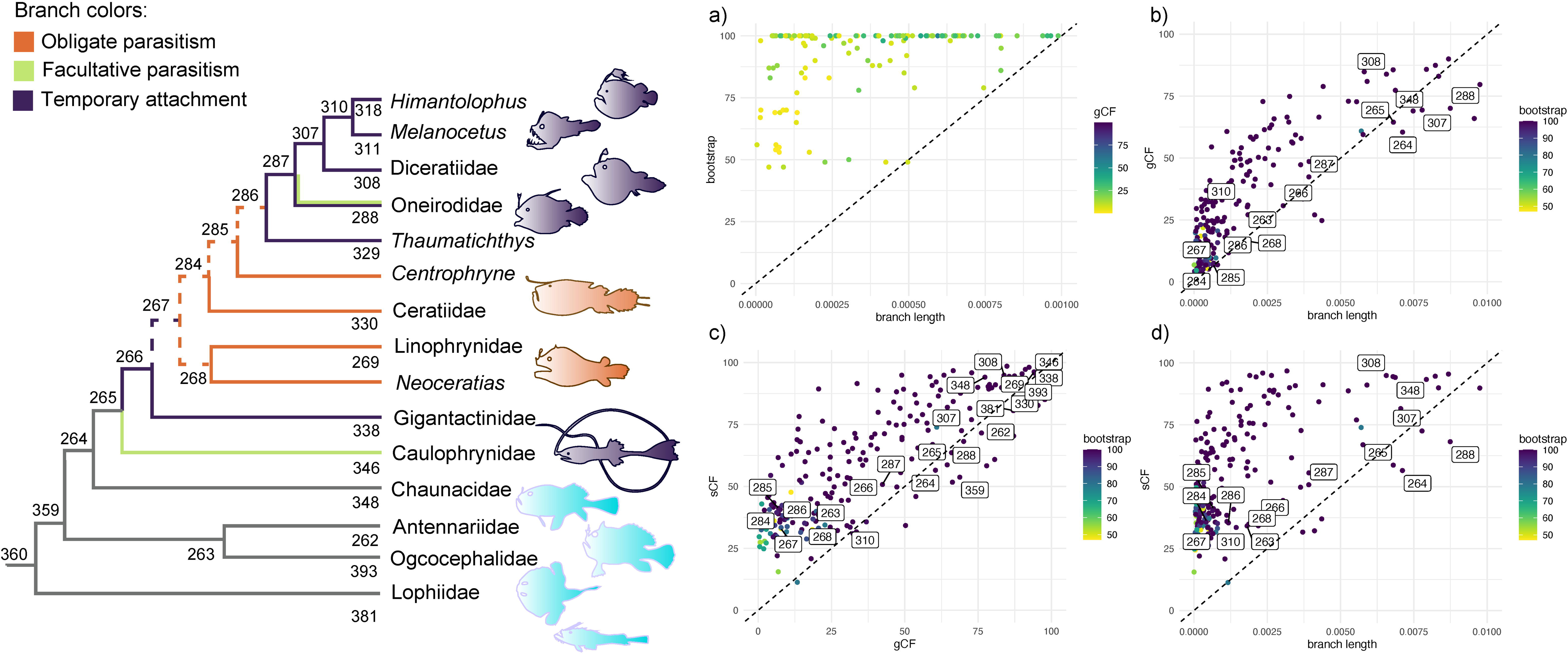
Sampling completeness. Map showing the coverage of the subset of UCEs selected following CIAlign.

**Figure S9.**
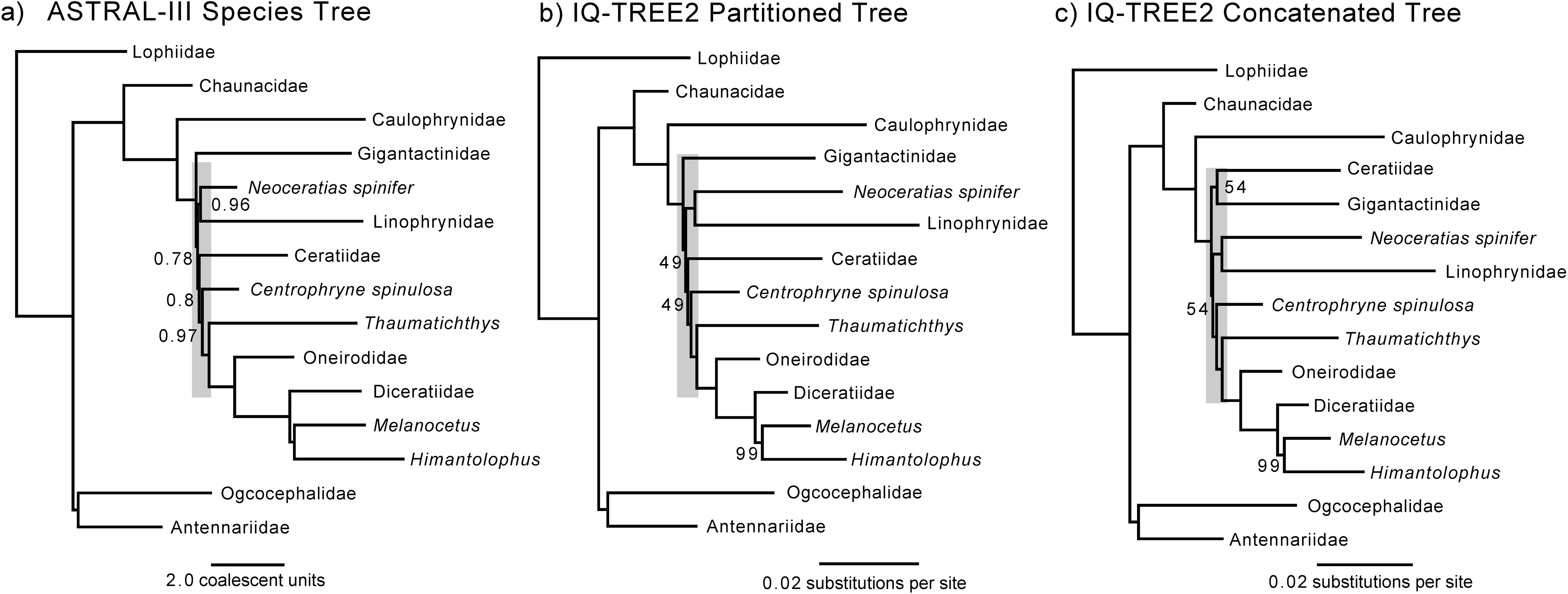
Concordance and discordance in anglerfish phylogeny. Scatterplots showing relationships between (a) bootstraps and branch lengths, (b) gene concordance factors and branch lengths, (c) site concordance factors and gene concordance factors, and (d) site concordance factors and branch lengths. Key nodes are numbered on the tree to the left.

**Figure S10.**
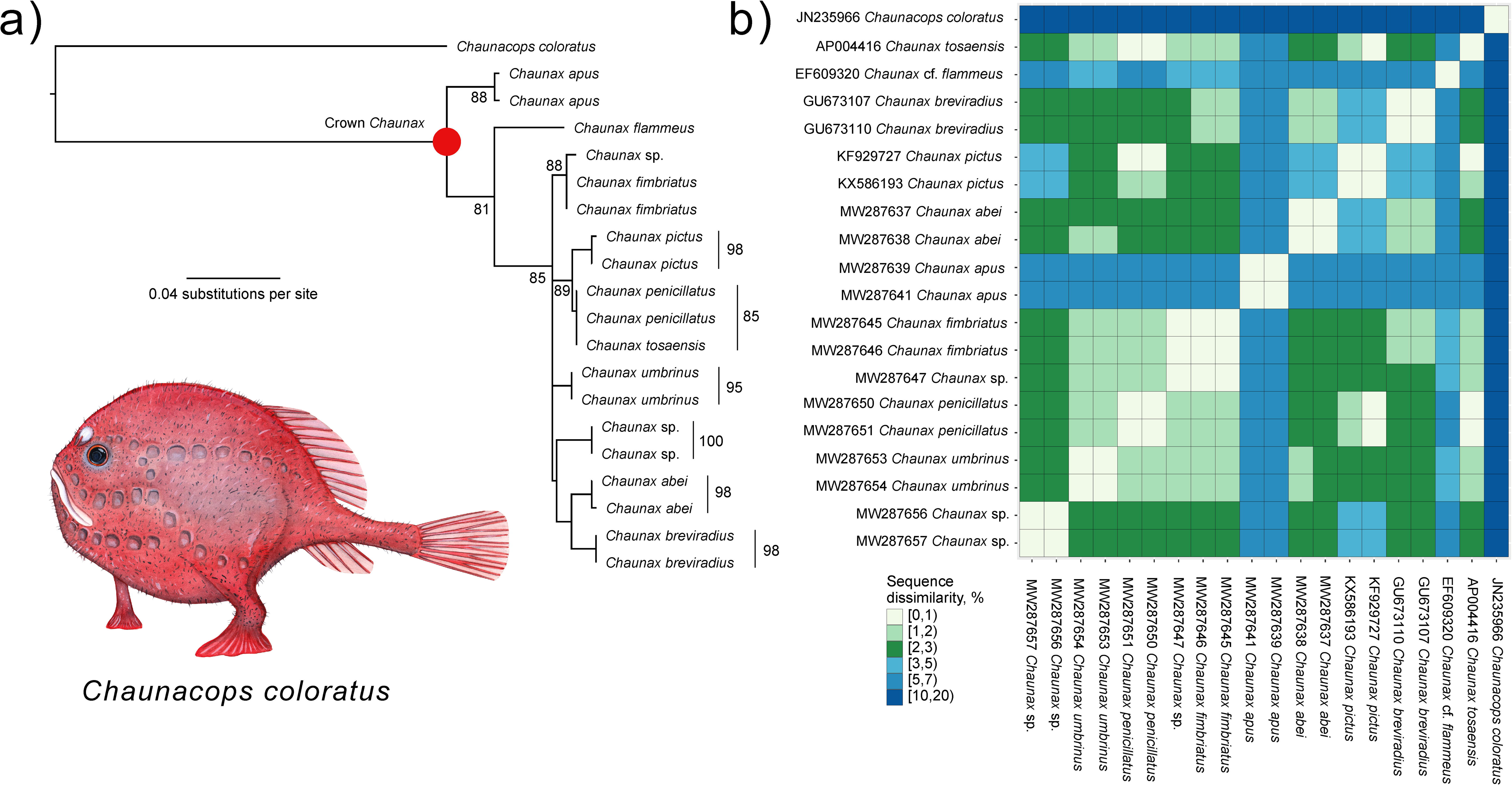
The anomaly zone in ceratioid phylogeny. Simplified topologies resulting from (a) the ASTRAL III species tree summary of individual gene trees generated in IQ-TREE2, (b) the maximum likelihood phylogeny generated from the analysis of partitioned UCE sets in IQ-TREE2, and (c) the analysis in IQ-TREE2 where all UCEs were concatenated. Shaded gray indicates the location of the detected anomaly zone in each phylogeny.

**Figure S11.**
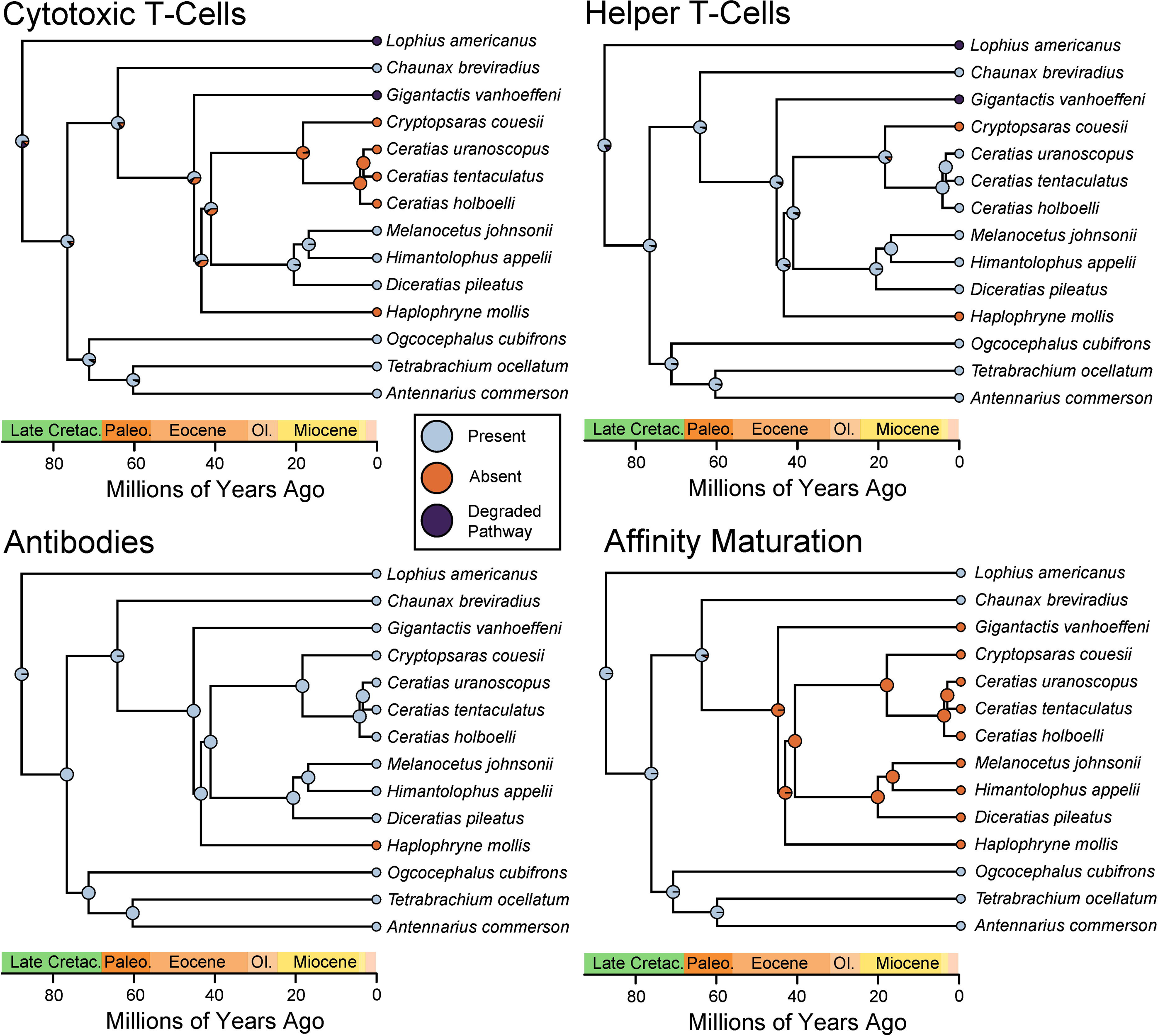
Chaunacid diversity. Maximum likelihood tree of the mitochondrial locus *COI* Showing relationships between species in *Chaunacidae*. Red dot indicates the crown node for *Chaunax*. (b) Heatmap showing uncorrected genetic dissimilarity values between species in Chaunacidae. Note the poor resolution within *Chaunax*. Illustration by Julie Johnson.

**Figure S12.**
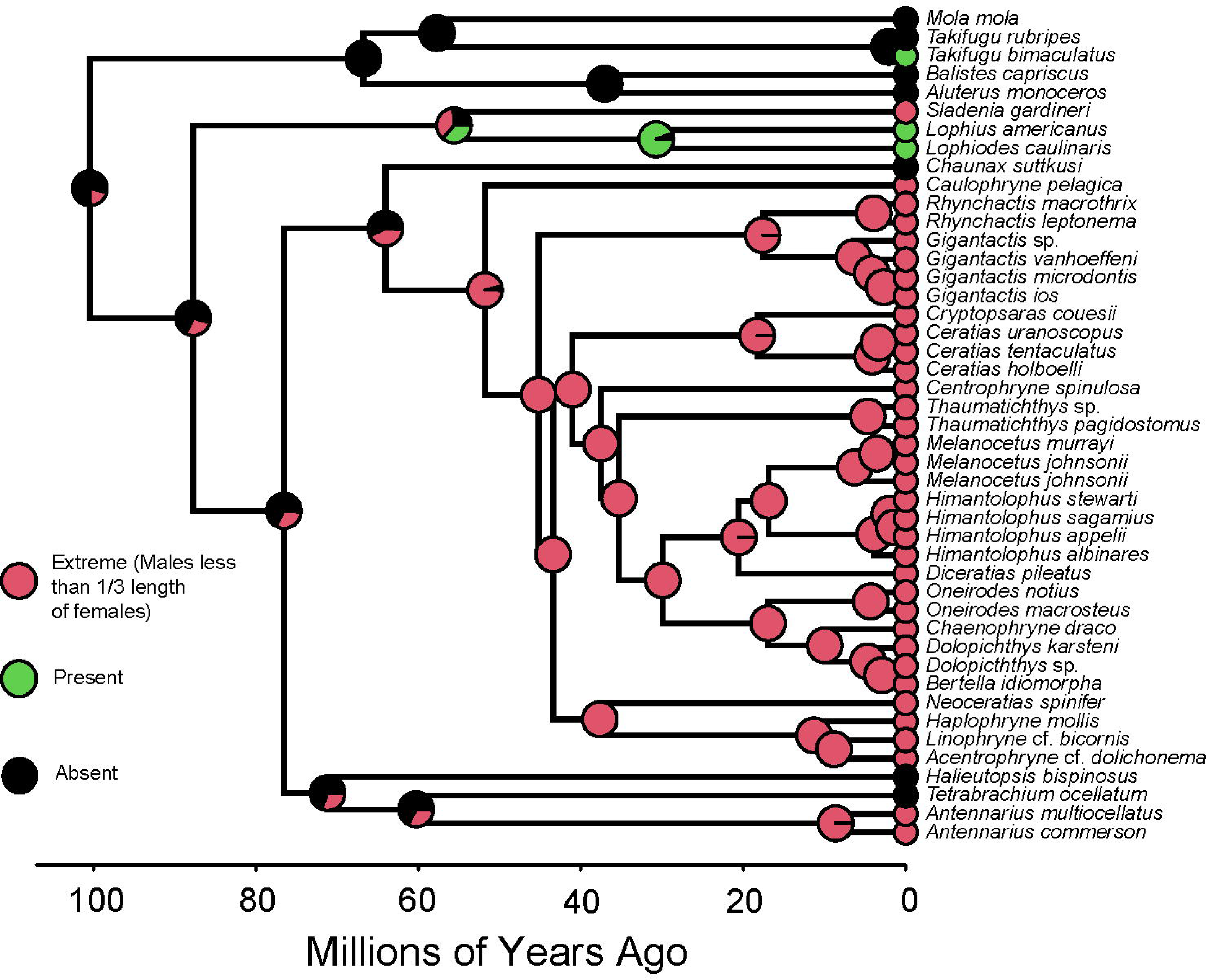
Convergent Degeneration of Adaptive Immunity in Anglerfishes. Simplified phylogeny with ancestral state reconstructions of different components of the vertebrate adaptive immune system using stochastic character mapping (R phytools) on a simplified version of the anglerfish timetree presented in Figure 1.

**Figure S13.**
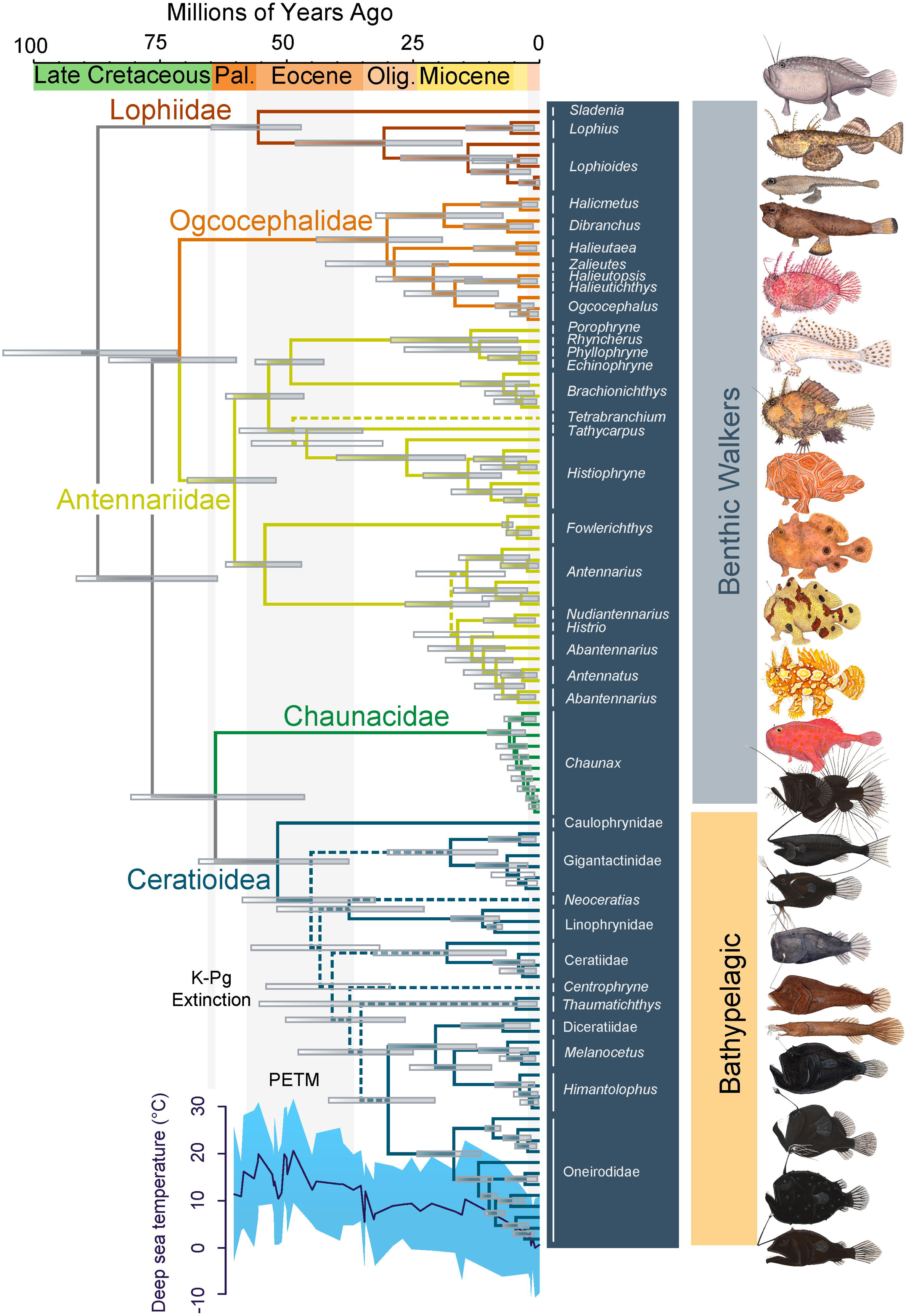
Evolution of Sexual Dimorphism in Anglerfishes. Time tree with ancestral state reconstructions of sexual dimorphism in anglerfishes using stochastic character mapping (R phytools) on a reduced version of the anglerfish timetree presented in Figure 1.

## Notes

### Competing Interest Statement

The authors have declared no competing interest.

